# Distinct Inductions of and Responses to Type I and Type III Interferons Promote Infections in Two SARS-CoV-2 Isolates

**DOI:** 10.1101/2020.04.30.071357

**Authors:** Fu Hsin, Tai-Ling Chao, Yun-Rui Chan, Han-Chieh Kao, Wang-Da Liu, Jann-Tay Wang, Yu-Hao Pang, Chih-Hui Lin, Ya-Min Tsai, Jing-Yi Lin, Sui-Yuan Chang, Helene Minyi Liu

**Affiliations:** Department and Graduate Institute of Biochemistry and Molecular Biology, No. 1 Sec. 1 Ren-Ai Road, Taipei City, Taiwan; Department of Clinical Laboratory Sciences and Medical Technology, College of Medicine, National Taiwan University, No. 1 Sec. 1 Ren-Ai Road, Taipei City, Taiwan; Department of Medicine, National Taiwan University Cancer Center, Taipei, Taiwan; Department of Internal Medicine, National Taiwan University Hospital, Taipei, Taiwan

## Abstract

The recent emerging coronavirus, SARS-CoV-2, has been rapidly and widely spread and causing an ongoing viral pneumonia outbreak worldwide. It has been observed that SARS-CoV-2 patients show a rather long and asymptomatic incubation time. We characterized the abilities to induce and to response to IFNβ/IFNλ1 of two or our clinical isolates, SARS-CoV-2/NTU01/TWN/human/2020 and SARS-CoV-2/NTU02/TWN/human/2020, which exhibit only two amino acid differences over the ∼30kb viral genome. We found that both isolates may infect Huh7, A549 and Calu-3 cells, yet the RIG-I-like receptor-dependent antiviral signaling was poorly induced in these cells in the early infections. Unexpectedly, we found that the intracellular vRNA levels of these isolates were sustained upon to type I/III IFN treatments, and this phenotype was more pronounced in the Taiwan/NTU01/2020 isolate. The type I/III IFN responses are antiviral but partially proviral in the case of SARS-CoV-2 infections. Poor induction and response to innate immunity may contribute to destitute neutralization index of the antibody produced, and indeed we found that the patient serum could not efficiently neutralize SARS-CoV-2 virions. With better understandings of the interplay between SARS-CoV-2 and the host antiviral innate immunity, our report may provide new insights for the regimen of therapies for SARS-CoV-2 infected patients.

## Main

In December 2019, a few pneumonia patients with low respiratory infections were identified in Wuhan, Hubei Province, China^1^. The etiology of this illness was then termed as coronavirus infection disease 2019 (COVID-19), which was caused by severe acute respiratory syndrome coronavirus 2 (SARS-CoV-2). SARS-CoV-2 shares about 80% nucleotide identity to the original SARS-CoV, and the corresponding disease, COVID-19, shows many similar symptoms to the original SARS-CoV infection^2^. Both SARS-CoV and SARS-CoV-2 can be transmitted from human to human, and both utilize human angiotensin converting enzyme 2 (ACE2) as the major receptor for virus entry^3-5^. However, the transmission of SARS-CoV-2 between human seems more efficiently than that of the original SARS-CoV. In April 2020, more than 2,000,000 cases of SARS-CoV-2-infected patients have been reported in more than 185 countries around the world.

The induction and expression of type I and type III interferons (IFNs) are the frontlines to fight against viral infections^6,7^. These responses occur within minutes to hours post-infection and rapidly set the infected and the neighboring cells in the anti-viral state by inducing the expression of interferon and antiviral immunity by recruiting innate and adaptive immune cells from hours to days post-infection. Upon virus infection and pathogen-associated molecular pattern (PAMP) recognition, the PAMP recognition receptors (PRRs) are then lead to the activation of PAMP-driven transcription factor, IFN production and interferon-stimulated gene (ISG) expression resulting in the immediate expression of host response genes^8^. An early onset of type I IFN induction and response is the key for successful viral clearance^7^. Particularly, type III IFN plays a critical role at the barrier surfaces, such as the airway and the GI tract^9^. It has been shown that IFNλ1 was more potent than type I IFN in restricting viral propagation with less inflammation and tissue damage ^10,11^. In contrast, many viruses have evolved some strategies to escape from the induction of response to type I/III IFN to establish the infection ^12,13^.

Asymptomatic SARS-CoV-2-infected individuals has been reported in many studies ^14^. Some SARS-CoV-2 infected patients showed long incubation time before the symptoms were developed. Two SARS-CoV-2-infected patients admitted to our unit showed distinct clinical virology and serology progresses (Table 1). Both patients showed normal complete blood count (CBC) initially with mild thrombocytopenia. Although days taken for seroconversion highly varied between the two patients, the immunoglobulin levels including total IgM, IgG and IgA were comparable (Table 1). While Patient NTU2 showed early seroconversion and rapid viral clearance, Patient NTU1 showed a slightly delayed seroconversion and a prolonged virus shedding period post seroconversion, up to 63 days post symptom onset (Table 1) ^15^. Therefore, it is likely that the anti-SARS-CoV-2 innate immunity was not sufficiently induced in time to confront SARS-CoV-2 infections in Patient NTU1. We then isolated viruses from both patients, SARS-CoV-2/NTU01/TWN/human/2020 (GeneBank: MT066175.1) and SARS-CoV-2/NTU02/TWN/human/2020 (GeneBank: MT066176.1), and assessed how these isolates may induce and response to host innate immunity in cell cultures. We then performed many assays in cell culture, where only the innate immunity of the infected host cells are involved, to rule out the variations among infected individuals and to characterize the virology of SARS-CoV-2 isolates. RNA sequencing results determined that SARS-CoV-2/NTU01/TWN/human/2020 represented clade B with the L84S (28144T>C) polymorphism in ORF8, and SARS-CoV-2/NTU02/TWN/human/2020 was highly related to the Wuhan prototype strain (Table 2). We hypothesize that the induction levels of type I and type III IFNs may be low during SARS-CoV-2 infection, and this may be due to the lack of PRR recognition and/or a strong suppression of type I/III IFN expressions by the viral proteins of SARS-CoV-2. To address these questions, we utilized two of our clinical isolates of SARS-CoV-2 to infect Huh7, A549, and Calu3 cells, and monitored their inductions and responses to type I/III interferons during SARS-CoV-2 infections.

**Table 1.** Summary of the clinical virology and serology findings in patients

| Patient | NTU1* | NTU2 |
| --- | --- | --- |
| Age | 50 | 45 |
| Sex | Female | Male |
| Initial CBC findings | mild thrombocytopenia (143 K/μL) | mild thrombocytopenia (129 K/μL) |
| Seroconversion | 10<sup>th</sup> day after symptom onset | 1<sup>st</sup> day after symptom onset |
| Immunoglobulin after seroconversion | IgG 1320 mg/dL | IgG 924 mg/dL |
|  | IgM 109 mg/dL | IgM 90.7 mg/dL |
|  | IgA 243 mg/dL | IgA 106 mg/dL |
| Virus sequence date | 9<sup>th</sup> day after symptom onset | 2<sup>nd</sup> day after symptom onset |
| Last date of virus isolation | 18<sup>th</sup> day after symptom onset | 5<sup>th</sup> day after symptom onset |
| Last date of vRNA detected | 63<sup>rd</sup> day after symptom onset | 15<sup>th</sup> day after symptom onset |
| CRP records | Peaked at 10<sup>th</sup> day | Back to normal levels on the 7<sup>th</sup> day |
| Virus isolate | SARS-CoV-2/NTU01/TWN/human/2020 (MT066175.1) | SARS-CoV-2/NTU01/TWN/human/2020 (MT066176.1) |

\*: data summarized from a previous report (Liu et.al., 2020).

**Table 2.**
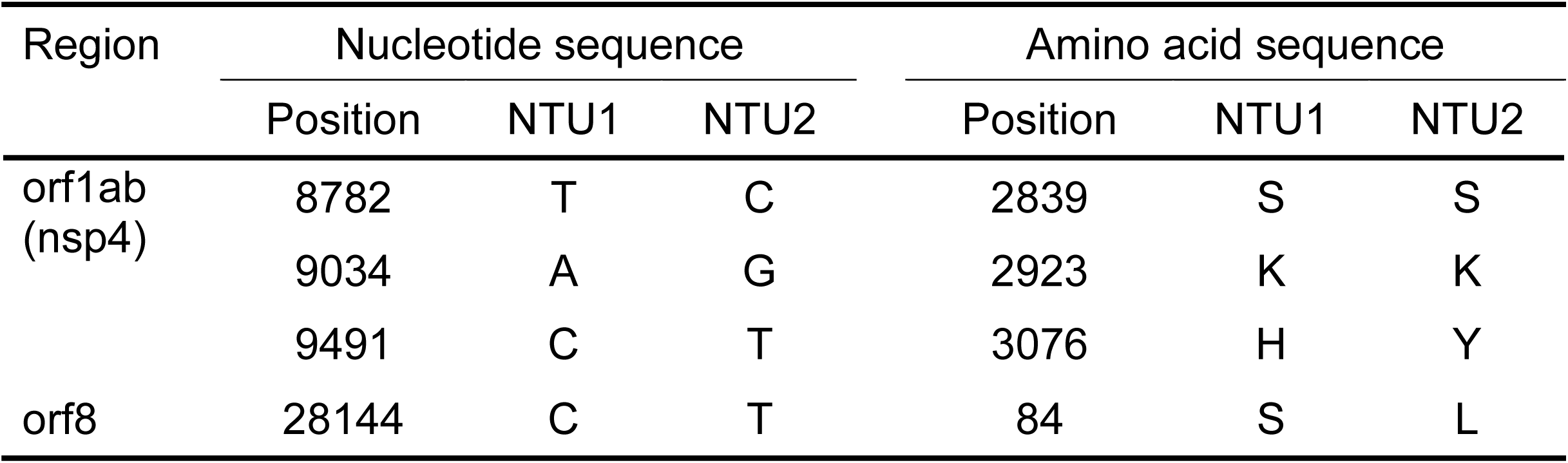
Sequence comparison of SARS-CoV-2/NTU01/TWN/human/2020 (NTU1) and SARS-CoV-2/NTU02/TWN/human/2020 (NTU2)

We first monitored the intracellular vRNA levels of SARS-CoV-2 in Huh7 hepatoma cells, of which cell-line that was widely used for previous SARS-CoV studies. Huh7 and MAVS K/D Huh7 cells were infected with either SARS-CoV-2/NTU01/TWN/human/2020 (NTU1) or SARS-CoV-2/NTU02/TWN/human/2020 (NTU2) at the multiplicity of infection (m.o.i.) of 0.01-0.1 for two days. By 48 hours post infection (h.p.i.), we did not observe any cytopathogenic effects (CPE) in Huh7 nor in the MAVS K/D huh7 cells (Supplementary Fig. 1A). The total intracellular vRNA was reverse-transcribed to cDNA, and the levels were determined by Taqman qPCR. As expected, the detected intracellular SARS-CoV2 vRNA levels increased along with the initial m.o.i. used for infection in both NTU1- and NTU2-infected Huh7 cells (Fig 1A). For both isolates, the vRNA levels was higher at 24 h.p.i than those at 48 h.p.i, suggesting that the replication of SARS-CoV-2 in Huh7 is none or very minimal (Fig. 1A). Also, NTU1 and NTU2 showed comparable intracellular vRNA levels in the infected Huh7 cells (Fig. 1A). Compared to the wildtype Huh7 cells, we did not observe any significant differences at the intracellular vRNA levels in the SARS-CoV-2 infected MAVS K/D huh7 cells (Fig. 1B). The loss of MAVS-dependent antiviral innate immunity did not further support replication of SARS-CoV-2 in Huh7 cells (Fig. 1B), suggesting that the RIG-I-like receptors-dependent antiviral innate signaling, primarily the induction of type I and type III IFNs, may only have marginal anti-viral effects on the replication of SARS-CoV-2. As an infection control of RNA viruses, Sendai virus (SenV), which belongs to the paramyxoviridae, was utilized. We then compared the induction levels of IFNβ and IFNλ1 in SARS-CoV-2-infected and SenV-infected Huh7 cells. After SenV-infection, Huh7 cells generated decent antiviral innate immunity against SenV as the mRNA levels of IFNβ, IFNλ1, and IFIT were all elevated (Supplementary Fig. 1B). Besides, the mRNA levels of IFITM3, which is specifically controlled by the interferon-sensitive response element (ISRE) downstream of type I IFN receptors, was also elevated, suggesting that Huh7 cells have the intact inductions and responses to type I IFN during SARS-CoV-2 infection (Supplementary Fig. 1B). However, in Huh7 cells infected with SARS-CoV-2 at 0.01 m.o.i., we found that IFNβ mRNA levels were only transiently induced with a very low fold of induction at 24 h.p.i and were down to background levels at 48 h.p.i in the infected Huh7 cells, which is correlating to the vRNA levels detected in these cells (Fig. 1, A & C). We did not observe any significant induction of IFNβ in Huh7 cells infected with SARS-CoV-2 at 0.1 m.o.i. (Fig. 1C).

**Figure 1.**
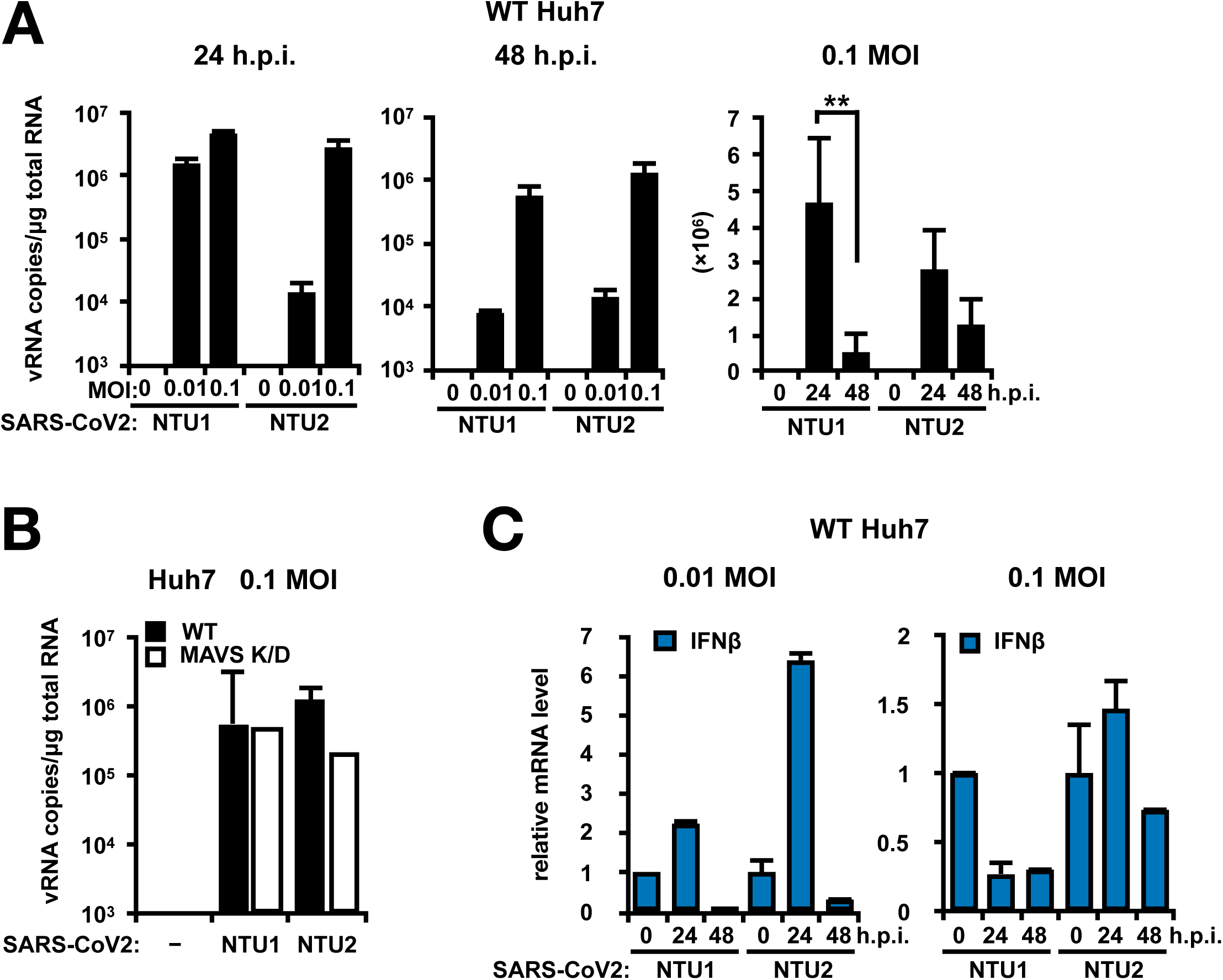
SARS-CoV2 vRNA levels and type I/III IFN mRNA levels in SARS-CoV2 infected Huh7 and MAVS-KD Huh7 cells. (A) The intracellular vRNA levels in SARS-CoV2 infected Huh7 cells. Huh7 cells were infected with either NTU1 or NTU2 SARS-Cov2 isolates at indicated m.o.i. for 24 or 48 hour prior to total RNA extraction. The vRNA levels of both isolates in SARS-CoV2 infected Huh7 cells decreased over time, especially for the NTU1 isolate. (B) SARS-CoV2 vRNA levels in either Huh7 or MAVS K/D Huh7 cells at 24 hours post-infection were at comparable levels. (C) The total RNA from (A) was examined for the mRNA expressions of type I IFN by qRT-PCR. IFNβ mRNA levels were transiently induced at very low levels in Huh7 cells infected with SARS-CoV2 at 0.01-0.1 m.o.i..

A recent study has reported that the virus-like particle of SARS-CoV-2 could enter Huh7, A549, Calu-3, and many other cell lines ^3^. We then repeated above experiments in relevant airway cell-lines, A549 and Calu-3 cells, which are adenocarcinomic human alveolar basal epithelial cells and human lung cancer epithelial cell, respectively. In consistent with data collected from Huh7 cells, both isolates were able to infect A549 and Calu-3 cells (Fig. 2, A & B). Similarly, the vRNA levels in A549 was higher at 24 h.p.i than those at 48 h.p.i (Fig. 2A), which is consistent to the phenotype observed in SARS-CoV-2 infected Huh7 cells, suggesting that A549 cells could not efficiently support SARS-CoV-2 replication either. Strikingly, Calu-3 cells were capable to highly efficiently support SARS-CoV-2 replication as shown by the increased intracellular vRNA levels over time from 24 to 48 h.p.i. with SARS-CoV-2 infection at 0.01 m.o.i (Fig. 2B). The vRNA levels in 0.1 m.o.i. SARS-CoV-2 infected Calu-3 cells were not examined due to the great loss of cells from cell death after 48 hours (data not shown). Among the three cell-lines that we infected with the same m.o.i, the intracellular vRNA levels in Calu-3 was 10,000-fold higher than those in the Huh7 and A549 cells (Figs. 1A & 2A-B), and Calu-3 is the only human cell-line we tested that showed a CPE after SARS-CoV-2 infection (Fig. 2C).

**Figure 2.**
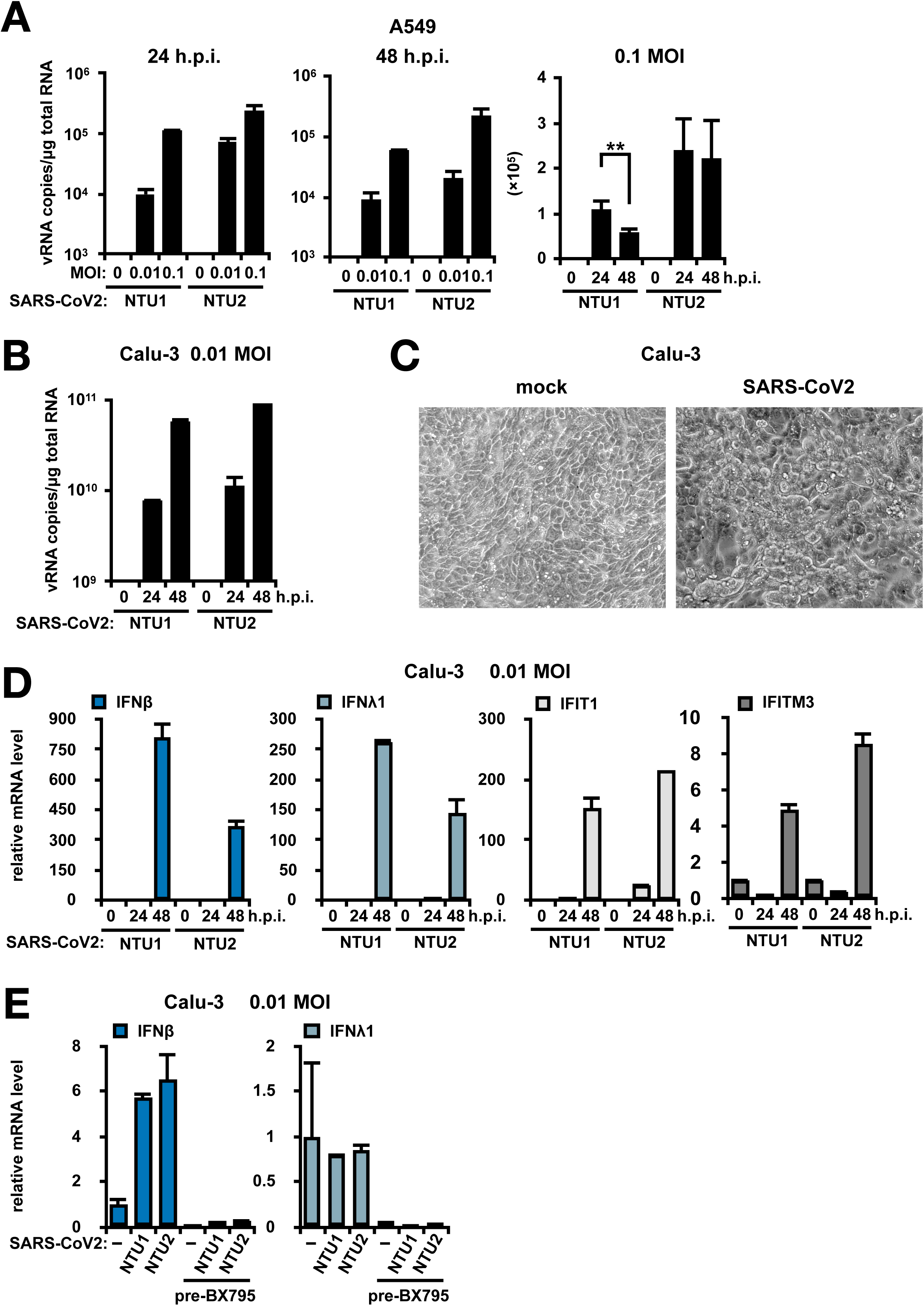
SARS-CoV2 vRNA levels and type I/III IFN mRNA levels in SARS-CoV2 infected A549 and Calu-3 cells. (A) The intracellular vRNA levels in SARS-CoV2 infected A549 cells. A549 cells were infected with either NTU1 or NTU2 SARS-Cov2 isolates at indicated m.o.i. for 24 or 48 hours prior to total RNA extraction. The vRNA levels of both isolates in SARS-CoV2 infected A547 cells were lower than those in Huh7 cells and also decreased over time. (B) Both SARS-CoV2 isolates efficiently infected and replicated in Calu-3 cells. The intracellular levels of vRNA increased overtime from 24-48 hours post-infection. (C) Cytopathogenic effect (CPE) of SARS-CoV2 (NTU1) infected Calu-3 cells. (D) Delayed induction of type I/III IFNs (IFNβ and IFNλ1) and ISGs (IFIT1 and IFITM3) expressions in SARS-CoV2 infected Calu-3 cells. The expression of all four target genes were nearly none at 24 h.p.i. and greatly induced after 48 h.p.i. despite the high vRNA levels at both 24 and 48 h.p.i as shown in (B). (E) The expression levels of IFNβ and IFNλ1 mRNAs in SARS-CoV2 infected Calu-3 cells were at low levels at 24 h.p.i., and this induction was inhibited by the pretreatment of 10 uM BX-795, which blocks the TBK1 kinase activity, prior to SARS-CoV2 infection.

Besides SARS-CoV-2 replication, the induction of type I/III IFNs in SARS-CoV-2 infected Calu-3 cells, which supports SARS-CoV-2 replication most efficiently among the cells we tested, were also assessed (Fig. 2D). To confirm that Calu-3 has intact type I/III IFN inductions and responses against RNA virus infections, we first infected Calu-3 cells with 100 HAU/ml SenV for 24-48 hours and examined the intracellular vRNA levels and the mRNA levels of ISGs (Supplementary Fig. 2A). As expected, SenV vRNA levels increased over time from 24 to 48 h.p.i. (Supplementary Fig. 2A), and the mRNA levels of IFNβ, IFNλ1, IFIT1, and IFITM3 were also increased, suggesting the SenV-infected Calu-3 was able to turn on the antiviral signaling against SenV infection. Conversely, although the intracellular levels of SARS-CoV-2 vRNA were readily amplified at 24 h.p.i, the induction of IFNβ and IFNλ1 mRNA expressions in Calu-3 remained undetectable at 24 h.p.i (Fig. 2, B & D), indicating either that the intracellular vRNAs of SARS-CoV-2 are poor ligands for RNA recognition by the PRRs to induce type I/III IFNs or certain viral products of SARS-CoV2 could strongly suppress the induction of antiviral innate immunity. This phenotype is very different from how Calu-3 response to SenV infection (Fig. 2D and Supplementary Fig. 2A). During SenV infection, the fold induction of IFNβ and IFNλ1 mRNA was correlated to the vRNA levels of SenV (Supplementary Fig. 2A). Nevertheless, during SARS-CoV-2 infection, the attitude of the induction of IFNβ and IFNλ1 mRNA expression was not correlated to the intracellular vRNA levels. The initial induction of IFNβ and IFNλ1 mRNA expression was weak at 24 h.p.i during SARS-CoV-2 infection (Fig. 2, D and E) At 48 hours post SARS-CoV-2 infection, along with the elevated amount of intracellular vRNA (Fig. 2B), both the mRNA levels of IFNβ and IFNλ1 in SARS-CoV-2 infected Calu-3 were drastically increased, and the expressions of ISGs, including IFIT and IFITM3 were also elevated (Fig. 2D). As previously mentioned that the expression of IFITM3 is under the control of ISRE, these results suggested that in Calu-3 cells, the production of type I/III IFNs was not impaired during SARS-CoV-2 infection (Fig. 2D). We next determined whether the induction of type I and type III IFNs were dependent on the TBK1-IRF3 axis of signaling. Calu-3 cells were infected with SARS-CoV-2 for 6 hours and then treated with a TBK1 inhibitor, BX795, for 18 hours. The IFNβ and IFNλ1 mRNA were induced at low levels after 24 hours SARS-CoV-2 infection in the mock-treated Calu-3 cells, and this induction was diminished in the BX-795 pre-treated SARS-CoV-2-infected Calu-3 cells (Fig. 2E), indicating the critical roles of TBK1-IRF3 in SARS-CoV-2-induced type I/III IFN induction. These data suggested that SARS-CoV-2 vRNA was only able to induce antiviral innate immunity in cells which well-supported SARS-CoV-2 replication, and that SARS-CoV-2 is likely a weak inducer and also a strong suppressor of type I/III IFNs.

Our results indicate that the viral RNA of SARS-CoV-2 was able to be detected by intracellular PRRs to induce type I/III IFN expression. Although the induction of IFNβ and IFNλ1 was transiently detected in SARS-CoV-2 infected Huh7 cells, the mRNA levels of several ISGs, including IFIT1 and IFMTF3 were not increased during infection (Fig. 3A). We then assessed whether the intracellular vRNA of SARS-CoV-2 could serve as the ligands of the cytoplasmic foreign RNA receptors, RIG-I and MDA5. Total RNA extracted from the NTU-1- or NTU-2-infected Calu-3 cells were transfected into mouse embryonic fibroblasts (MEFs) treated with cycloheximide to monitor their abilities to induce type I IFN expression by measuring the mRNA levels of *Ifnb* in the absence of any newly-translated proteins (Fig. 3B). As negative and positive controls, total Calu-3 intracellular from mock-infected (mcRNA) or Sendai virus-infected (SenV-icRNA) cells were also transfected into MEFs respectively (Fig. 3B). The transfection of mcRNA into MEFs did not induce any expression of *Ifnb* in MEFs. The *Ifnb* mRNA levels were greatly increased in MEFs transfected with total intracellular RNA harvested from SenV-infected cells (Fig. 3B). We then pretreated the SenV-icRNA samples with calf intestinal alkaline phosphatase prior to transfection to reduce RIG-I recognition of SenV-icRNA, and as expected, the *Ifnb* mRNA levels induced by CIP-treated SenV-icRNA transfection was drastically reduced (Fig. 3B). We found that MEFs transfected with the total RNAs extracted from SARS-CoV-2-infected Calu-3 cells (NTU1- and NTU2-icRNAs) were able to induce slight expression of *Ifnb* mRNA when compared to that transfected with SenV-icRNA (Fig. 3B), and the phosphatase pretreatment of the total RNA extracted from SARS-CoV-2-infected cells prior to RNA transfection could barely reduce the induction of *Ifnb* mRNA, suggesting that the SARS-CoV-2 RNA may be recognized mostly through PRRs other than RIG-I (Fig. 3B).

**Figure 3.**
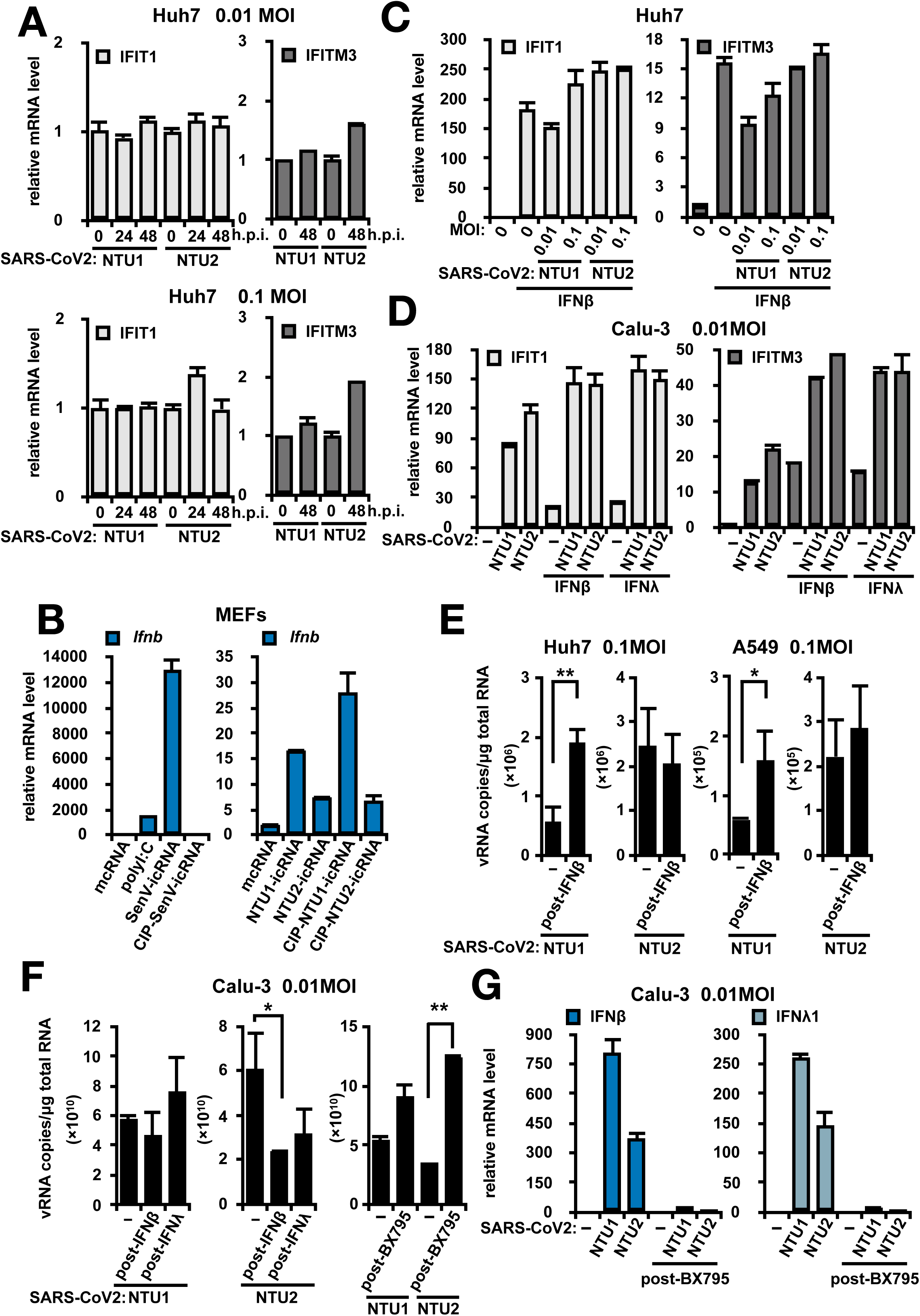
The induction of and response to Type I/III IFNs in SARS-CoV2 infected cells could not restrict the viral replication. (A) The expression of IFIT1 and IFITM in SARS-CoV2 infected Huh7 cells. The induction of expression of both IFIT1 and IFITM3 were weak or none. (B) Total RNA from SenV-, SARS-CoV NTU1- and NTU2-infected Calu-3 were extracted and transfected into cyclohexamine-treated mouse embryonic fibroblast to determine their ligand activities in activation RLR-dependent type I IFN induction. Mock-infected total RNA (mcRNA) was utilized as a negative control. NTU1-icRNA and NTU2-icRNA, which were extracted from the SARS-CoV2-infected Calu-3 cells could only moderately induce *Ifnb* expression in MEFs when compare to the RNA extracted from SenV-infected cells (SenV-icRNA). Phosphatase pretreatment by calf intestine phosphatase (CIP) could partially reduce the post-transfection induction of *Ifnb.* (C) Huh7 cells were infected with SARS-CoV2 for 24 hours and then treated with 100 IU/mL IFNβ for additional 24 hours. The expressions of IFIT1 and IFITM3 mRNAs in Huh7 cells were drastically increased under the treatment of IFNβ, and this response was not impaired by the infection of SARS-CoV2. (D) Calu-3 cells were infected with SARS-CoV2 for 24 hours and then treated with 100 IU/mL IFNβ or 100ng/mL IFNλ1 for additional 24 hours. The expressions of IFIT1 and IFITM3 mRNAs in Calu-3 cells were drastically increased under the treatment of IFNβ and/or IFNλ1, and this response was not impaired by the infection of SARS-CoV2. (E) The effects of post-infection treatment of IFNβ in Huh7 and/or A549 cells infected with SARS-CoV2. Post-treatment of IFNβ in Huh7 and/or A549 cell could not reduce the intracellular vRNA levels of SARS-CoV2, and the vRNA levels of NTU1 isolate in both Huh7 and A549 cells were increased upon IFNβ treatment. (F) The effects of post-infection treatment of IFNβ, IFNλ1, and BX795 in Calu-3 cells infected with SARS-CoV2. Calu-3 cells were first infected with SARS-CoV2 for 24 hours and then treated with indicated agents. Treatment of IFNβ and IFNλ1 post-infection could not restrict the intracellular vRNA levels of NTU1 isolate and was able to slightly reduce the intracellular vRNA levels of NTU2. The treatment of BX795 increased the intracellular vRNA level, especially that of the NTU2 isolate. (G) Treatment of BX795 in SARS-CoV2 cells drastically reduced both IFNβ and IFNλ1 mRNA expression induced by SARS-CoV2 infection.

Based on these results, it is suggested that although the replication intermediates of SARS-CoV-2 is an inducer for type I IFN expression, SARS-CoV-2 may encode some viral products as strong type I IFN induction suppressors, which together contribute the poor induction of type I/III IFNs (Figs. 2D and 3B). Therefore, we next assessed whether the complementarity by exogenous source of type I or type III IFNs in the culture medium may facilitate the cells to establish the antiviral states to restrict SARS-CoV-2 replication. Huh7 and Calu-3 cells were infected with SARS-CoV-2 isolates at the indicated m.o.i. for 24 hours and then treated with either 100 IU/mL of IFNβ or 100 ng/ mL of IFNλ1 for 24 hours. The intracellular RNA from these cells were then extracted by trizol and reverse-transcribed into cDNA for quantitative PCR. The mRNA levels of several ISGs, including IFIT1 and IFITM3, were first determined to validate that the treatment of IFNβ and/or IFNλ1 indeed turned on the expression of ISGs (Fig. 3C). The expression levels of ISGs post-interferon treatments were at comparable levels among the mock-, NTU1-, and NTU2-infected cells, suggesting that SARS-CoV-2 infection did not block the response to type I IFN in Huh7 cells (Fig. 3C). In Calu-3 cells, SARS-CoV-2 infection may induce the expression of type I/III expression at 48 h.p.i. (Fig. 2D), and responses to both type I and type III IFN post-treatments during SARS-CoV-2 further elevated the mRNA expression levels of IFIT1 and IFITM3 (Fig. 3D). Typically, the expression of ISGs in response to the treatment of type I/III IFNs facilitate the cells to be antiviral as we have shown that either pre- or post-treated the cells with type I/III IFN during SenV infection, the cells were able to restrict SenV replication (Supplementary Fig. 2, A & B). Calu-3 cells pretreated with type I and/or type III IFNs were also able to restrict SARS-CoV-2 replication (Supplementary Fig. 3A). However, in all cells which were post-treated with either type I and/or type III IFNs after SARS-CoV-2 infection, we did not observe any antiviral effects in these cells as the vRNA levels sustained after the treatment of type I/III IFNs (Fig. 3E). The vRNA levels of NTU2 remained unchanged upon type I IFN treatment in Huh7 and A549 cells (Fig. 3E), but however, the vRNA levels of NTU1 were predominantly enhanced after the treatment of type I IFN (Fig. 3E). In Calu-3 cells, which highly supports SARS-CoV-2 replication, the vRNA levels of NTU1 were not reduced upon post-treatment of type I/III IFNs while the vRNA levels of NTU2 were slightly suppressed by type I/III IFN treatments (Fig. 3F). In contrast, the post-treatment of BX795, which inhibits the expression of type I/III IFN (Fig. 3G), was able to increase the vRNA levels of NTU2 in Calu-3 cells, of which viral infection is more sensitive the type I/III IFN post-treatment (Fig. 3F). All these data indicated that pre-treatment of type I/III IFNs potentiates the cells to restrict SARS-CoV-2 infection, which is consistent to a recent report in SARS-CoV-2 infected Vero-E6 cells ^16^. However, once the infection of SARS-CoV-2 is established, the post-treatment of type I and/or type III IFNs could not effectively control SARS-CoV-2 replication. A recent report revealed the viral protein interactome of SARS-CoV-2 and showed that SARS-CoV-2 nsp13 was capable to interact with TBK1, which is a critical kinase to activation IRF3 for type I/III IFN expressions ^17^. The detail mechanism remained to be further studied.

It has been shown that deficiency in RLR-dependent antiviral signaling, such as the induction of type I IFN, may result in uncontrolled inflammation and adaptive immunity which could not provide protection against the viral infection^18^. The induction of type I IFN in the epithelial cells correlates to the neutralization index of antibodies produced during viral infection^18^. Our cell culture experiments showed that the inductions of type I/III IFN expressions were poor during SARS-CoV-2 infection, and therefore we next assessed whether the serum from SARS-CoV-2-infected patient may provide good quantity and quality of antibodies to neutralize SARS-CoV-2 virion. A recent study showed that among the hospitalized COVID19 patients, the occurrence of seroconversion was not followed by a rapid decline of the viral titer ^19^. For NTU1 patient, gradual viral load decrease was observed since hospitalization and the antibody response to SARS-CoV-2 was first identified on the tenth day ^15^. The plaque reduction neutralization test (PRNT) was performed to determine the neutralizing activities of serum collected from NTU2 patient post seroconversion and that from NTU1 patient on the 1^st^ and the 18^th^ day of hospitalization, which represented serum before and after seroconversion respectively^15^. The serum from NTU2 patient, who rapid clear SARS-CoV-2 after seroconversion (Table 1), was able to effectively neutralize NTU1 virions at >1:160 dilution in the PRNT assay (Supplementary Fig. 4). However, both serum collected from Patient NTU1 before and after seroconversion did not well-neutralize the virion isolated from the same patient (NTU1) The PRNT50 for NTU1 is about 1:80 before seroconversion and 1:160 after seroconversion (Fig. 4). These results together indicated that although the total IgM, IgG, and IgA amount between both patients were at the comparable or ever higher levels (Table 1), post-seroconversion serum from Patient NTU1 did not provide better neutralizations or protectivities against SARS-CoV-2 virions then the pre-seroconversion one.

**Figure 4.**
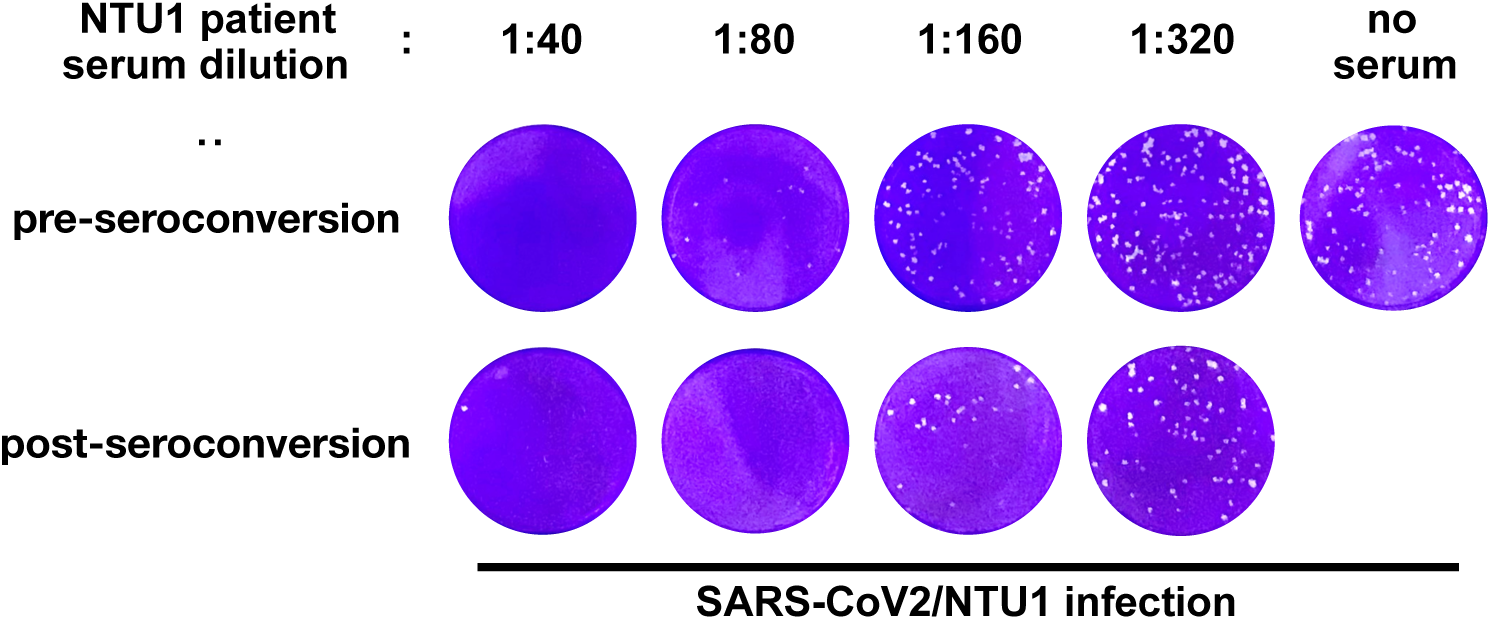
Plaque reduction neutralization of NTU1 isolate by NTU1 patient serum collected pre- and post-seroconversion.

In the early stage of the infection, the kinetics of virus replication and that of the establishment of antiviral state determines the infection outcome. With the ongoing outbreak of SARS-CoV-2 infection, characterizations of how SARS-CoV-2 replicates in the cell as well as how it may escape from the frontline of host antiviral innate immunity are critical for the development of therapy in responding to the novel infectious agents. During the preparation of this manuscript, several studies also reported that unlike SARS-CoV and/or Influenza A Virus infection, SARS-CoV-2 infection showed a delayed or suppressed phenotypes in the inductions of type I IFN and inflammatory cytokines in several models ^20,21^. However, in these experimental models, SARS-CoV-2 RNA replication was not ideal. We have tested several cell types and found that Calu-3, which supported SARS-CoV-2 replication with a CPE post-infection, may be a decent model to study this emerging virus instead of utilizing Huh7 and/or A549. Information obtained from SARS-CoV-2 infected Calu-3 cells will be relevant to the real infection situations. Our data suggested that although the vRNA and/or RNA replication intermediates could be recognized by host PRRs to induce type I/III IFN expressions, the virus likely encodes certain viral product to antagonize the antiviral innate immunity at the early stage of the infection. In some of our cull culture experiments, the treatment of type I or type III IFNs unexpectedly enhanced intracellular vRNA levels (Fig 3, E-F). Intriguingly, a recent report showed that among the top 50 genes which were co-expressed with ACE2, an entry receptor for SARS-CoV2, many of them were ISGs or genes that are involved in the regulation innate immunity, including ASS1, OAS1, MX1, and etc.^22^. In our study, we found that pretreatment of TBK1 inhibitor prior go SARS-CoV-2 infection, however, unexpectedly reduced the intracellular vRNA at 24 h.p.i., of which phenomenon also observed in the SARS-CoV-2-infected MAVS K/D Huh7 cells (Supplementary Fig. 3B and Fig. 1B). We examined the mRNA levels of ACE2 in our samples and found that the expression of ACE2 was increased after of IFNβ and/or IFNλ1 treatment and was decreased in cells treated with BX-795 or knocked-down with MAVS (Supplementary Fig. 3), which may be correlating with the reduced vRNA levels we observed in the MAVS K/D and BX795-treated cells during SARS-CoV-2 infection. A recent report showed that ACE2 may be an ISG in human epithelial cells ^23^. Our study together with recent reports suggest that SARS-CoV-2 may utilize certain ISGs to benefit its own life cycle. This may also explain why patients with chronic inflammation disease are more prone to COVID19. Whether the response to type I/III interferon treatments were purely anti-SARS-CoV-2 remained debatable.

The two isolates in this study, SARS-CoV-2/NTU01/TWN/human/2020 (NTU1) and SARS-CoV-2/NTU02/TWN/human/2020 (NTU2), presents similar replication preferences among the cell-lines we tested. The replication kinetics of these 2 isolates were similar, but however, their responses to the post-treatment of type I/III IFNs were different. While NTU2 was rather sensitive to the post-treatment of type I/III IFNs in the Calu-3 cells, NTU1 seemed to be resistant to the treatment of type I/III IFNs. The genetic difference of these 2 isolates were quite little. At the genetic level, there were positions 8782, 9034, 9491, and 28144 four nucleotides different. In the viral protein sequence, only two amino acid sites of orf1ab and orf8 were different (Table 2). Tang’s group found genetic analysis of 103 SARS-CoV-2 genomes by SNP could define the L and S lineages of SARS-CoV-2 in two linked SNPs at sites 8782 and 288144 ^24^. However, the virulence or pathogenicity of these two lineages of SARS-CoV-2 is still unclear. We tried to subclone and express ORF8 from both isolates in either mammalian cells or in E. coli, and none of the system could support decent expression of ORF8 (data not shown). Therefore, at this point, we are not able to design any molecular biological studies to elucidate whether the genotypic difference in ORF8 between the two isolates might contribute to the phenotypic differences in the response to type I/III IFNs.

Lastly, we found that seroconversion in patients did not correlate with their abilities for viral clearance. As reviewed a recent perspective, How the responses to type I/III IFNs in patients and/or healthy individual who received the vaccine may likely affect the protectivities provided by the antibodies^25^. Better assays to determine whether the antibodies developed during seroconversions are protective or pathogenic remained to be established. This will be rather useful to determine the regiment for each COVID19 patients as well as to evaluate the vaccine protectivity in the future.

## Material and Method

### Cells and Viruses

Huh7, A549, Calu-3, VeroE6 and LLC-MK2 cells were propagated in Dulbecco’s modified Eagle’s medium DMEM; Gibco-BRL, USA) supplemented with 10% fetal bovine serum (FBS; GIBCO-BRL, USA), penicillin G sodium 100 units/mL, streptomycin sulfate 100 μg/mL and amphotericin B 250 ng/mL (antibiotic-antimycotic; Gibco-BRL, USA). Stable MAVS knock-down (MAVS K/D) Huh7 cells as previously described^26^ was cultured in DMEM with 10% FBS and 1μg/mL of puromycin. Throat-swab, sputum, and/or nasopharyngeal swab specimens obtained from SARS-CoV-2-infected patient were maintained in viral-transport medium. The specimens were propagated in VeroE6 cells and/or LLC-MK2 cells in DMEM supplemented with 2 μg/mL of tosylsulfonyl phenylalantyl chloromethyl ketone (TPCK) -trypsin (Sigma-Aldrich). Culture supernatants were harvested when more than 70% of cells showing cytopathic effects, and the virus titers were determined by plaque assay. Experiments involving infectious SARS-CoV-2 followed the approved standard operating procedures of our Biosafety Level 3 facility at the College of Medicine, National Taiwan University.

### Virus infection

Virus infection was performed in 24-well tissue culture plates. The VeroE6 cells were seeded at 2 × 10^5^ cells/well in DMEM with 10% FCS and antibiotics one day before infection. SARS-CoV-2 at indicated MOI were added to the cell monolayer for 1 hour at 37°C. Subsequently, viruses were removed and the cell monolayer was washed once with PBS before covering with media containing 2% FCS. At the end of infection, the virus titers in the supernatants were determined by RT-qPCR (or plaque assay).

### RNA extraction and Quantitative reverse-transcription PCR

Total RNA from infected cells was extracted with Trizol reagent (Life Technologies) by isopropanol precipitations, and cDNA was prepared with iScript (BioRad). Quantitative PCR of vRNA was performed with TaqMan probes detecting SARS-CoV-2 N gene (IDT, #10006600). RP was used as a reference gene. Quantitative PCR of IFN-beta, IFN-lambda, IFIT1, and IFITM3 were performed with Power SYBR Green PCR (Life Technologies). Data were analyzed according to the ΔCT method. Primer and probe sequences are listed in Supplementary Table 1.

### RNA transfection

RNA transfection assay was described exhaustively in previous study^27^. Briefly, total RNA from calu-3 cells infected with SeV, SARS-CoV-2/NTU1 or SARS-CoV-2/NTU2 and uninfected cells were extracted with Trizol. All RNA samples were treated with or without Alkaline Phosphatase, Calf Intestinal (CIP) (New England BioLabs) for 30 min and extracted by Ethanol precipitations. Mouse embryonic fibroblasts (MEFs) were pre-treated with 20μg/mL cycloheximide for 30 min and then transfected with prepared RNA or polyI:C by TransIT-mRNA transfection kit (Mirus Bio) following manufacturer’s protocol. 8 hours post-transfection, total RNA was extracted with Trizol and analyzed by qPCR.

### Plaque reduction neutralization assay

Plaque reduction neutralization (PRNT) assay was performed to determine the SARS-CoV-2 titers as previously described with minor modification ^28^. Briefly, the VeroE6 cells were seeded at 2 × 10^5^ cells/well in the 24-well tissue culture plates with DMEM containing 10% FCS and antibiotics one day before infection. Serial diluted serum was preinteracted with SARS-CoV-2 at 37°C for one hour and then added to the cell monolayer for 1 hour at 37°C. Subsequently, viruses were removed, and the cell monolayer was washed once with PBS before covering with media containing 1% methylcellulose for 5-7 days. The cells were fixed with 10% formaldehyde overnight. After removal of overlay media, the cells were stained with 0.7% crystal violet and the plaques were counted.

### Statistic analysis

All data were presented the means ± standard deviation (SD) from two independent experiments, and the two-tailed Student’s t-test was used as statistical analysis, *p<0.05, **p<0.01.

## Ethical approval statement

The study was approved by the Research Ethics Committee or Institutional Review Board of the NTUH Research Ethics Committee (202002002RIND) and informed consent was waived.

## Acknowledgements

The authors would like to thank lab members in the Liu Lab and in the Chang Lab for their technical support. We thank Dr. Yu-Huan Tsai (Yang-Ming University), Dr. Hao-Sen Chiang and Dr. Li-Chung Hsu (National Taiwan University) for the insightful discussions. We thank the staff of the Biomedical Resource Core at the First Core Labs, National Taiwan University College of Medicine, for technical assistance. This research was supported by MOST (Ministry of Science and Technology, R.O.C, https://www.most.gov.tw/?l=en) grant MOST 108-2320-B-002-026-MY3, MOST 108-2320-B-002-049 and MOST107-2320-B-002-016-MY3.

## Author contributions

F.H and Y.R.C performed molecular studies of the gene expressions. T.L.C., H.C.K, Y.H.P, C.H.L., and Y.M.T. performed SARS-CoV2 infection experiments. H.M.L wrote the draft of the manuscript. J.Y.L, S.Y.C., and H.M.L edited the manuscript. S.Y.C. and H.M.L. conducted the research study.

## Supplementary Figure Legend

**Supplementary Fig. 1.**
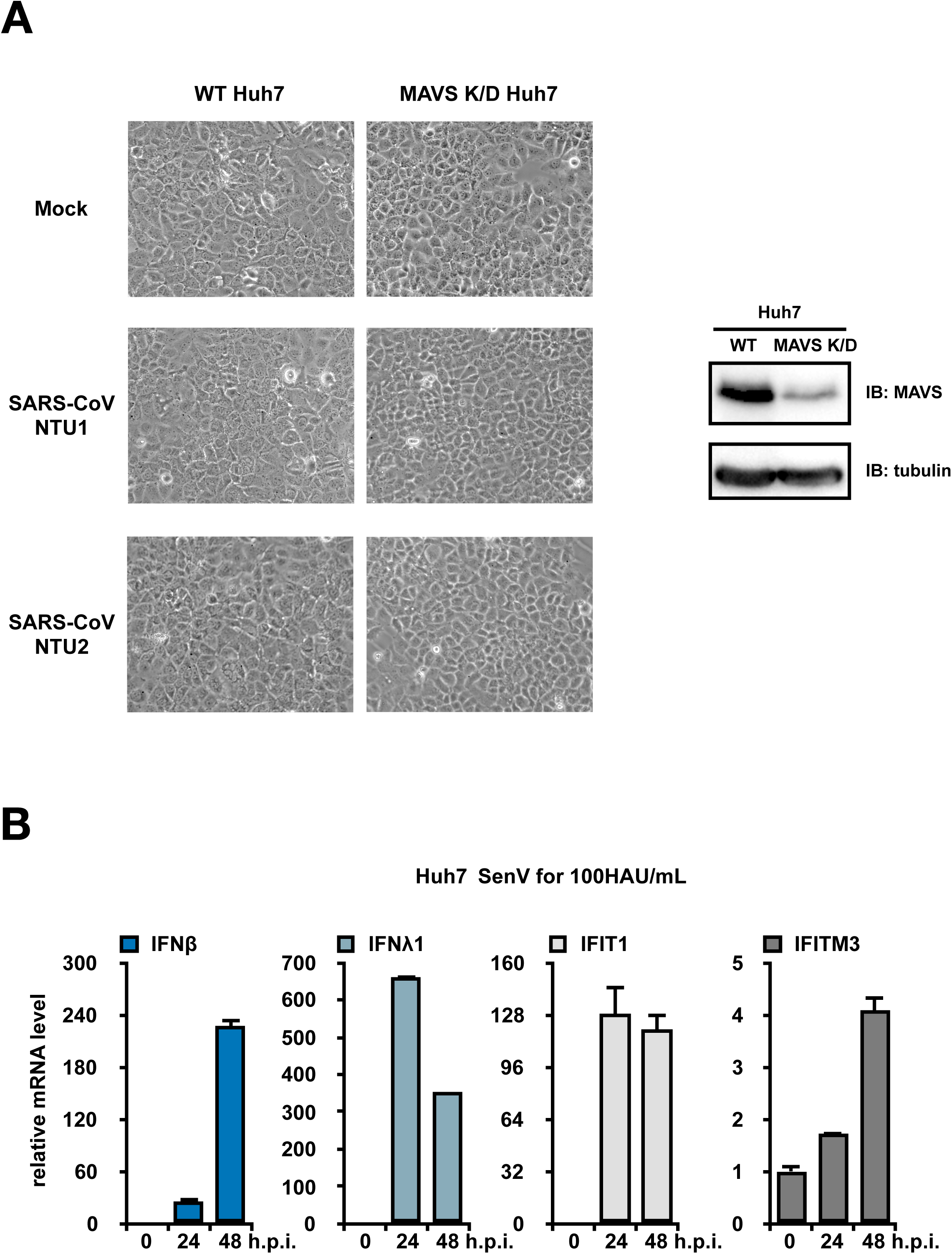
(A) SARS-CoV2 infected Huh7 and/or MAVS K/D Huh7 showed no CPE after infection. The protein expression levels of MAVS in both Huh7 and MAVS K/D Huh7 cells were determined by immunoblotting. (B) The expression of IFNβ, IFNλ1, IFIT1, and IFITM3 mRNA were all elevated in Huh7 cells in response to SenV infection.

**Supplementary Fig. 2.**
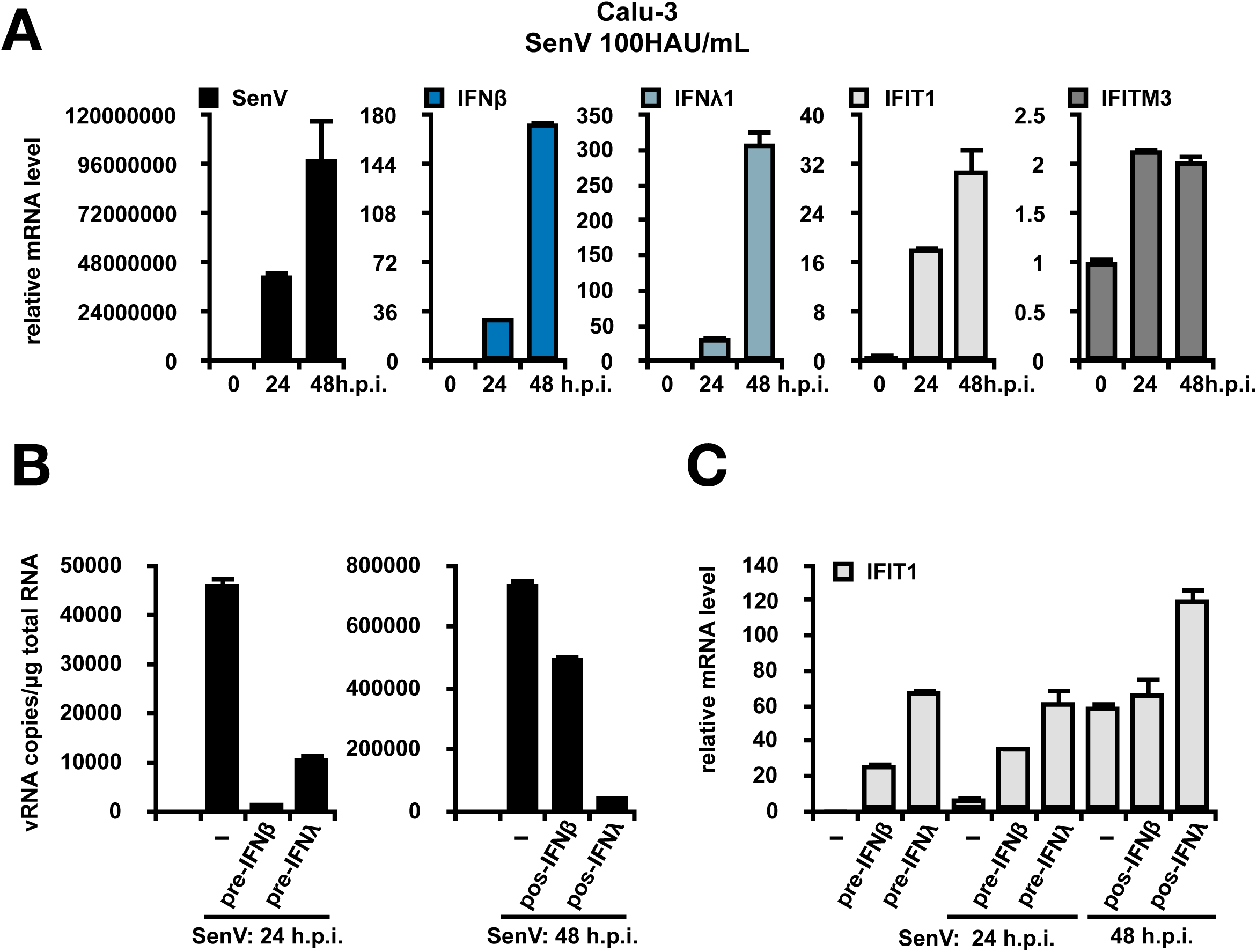
(A) The expression of IFNβ, IFNλ1, IFIT1, and IFITM3 mRNA were all elevated in Calu-3 cells inresponse to SenV infection. (B) SenV viral RNA was greatly reduced by either pre- and/or post-treatments of IFNβ and/or IFNλ1. (C) The mRNA levels of IFIT1 under the treatment of IFNβ and/or IFNλ1 and the infection of SenV.

**Supplementary Fig. 3.**
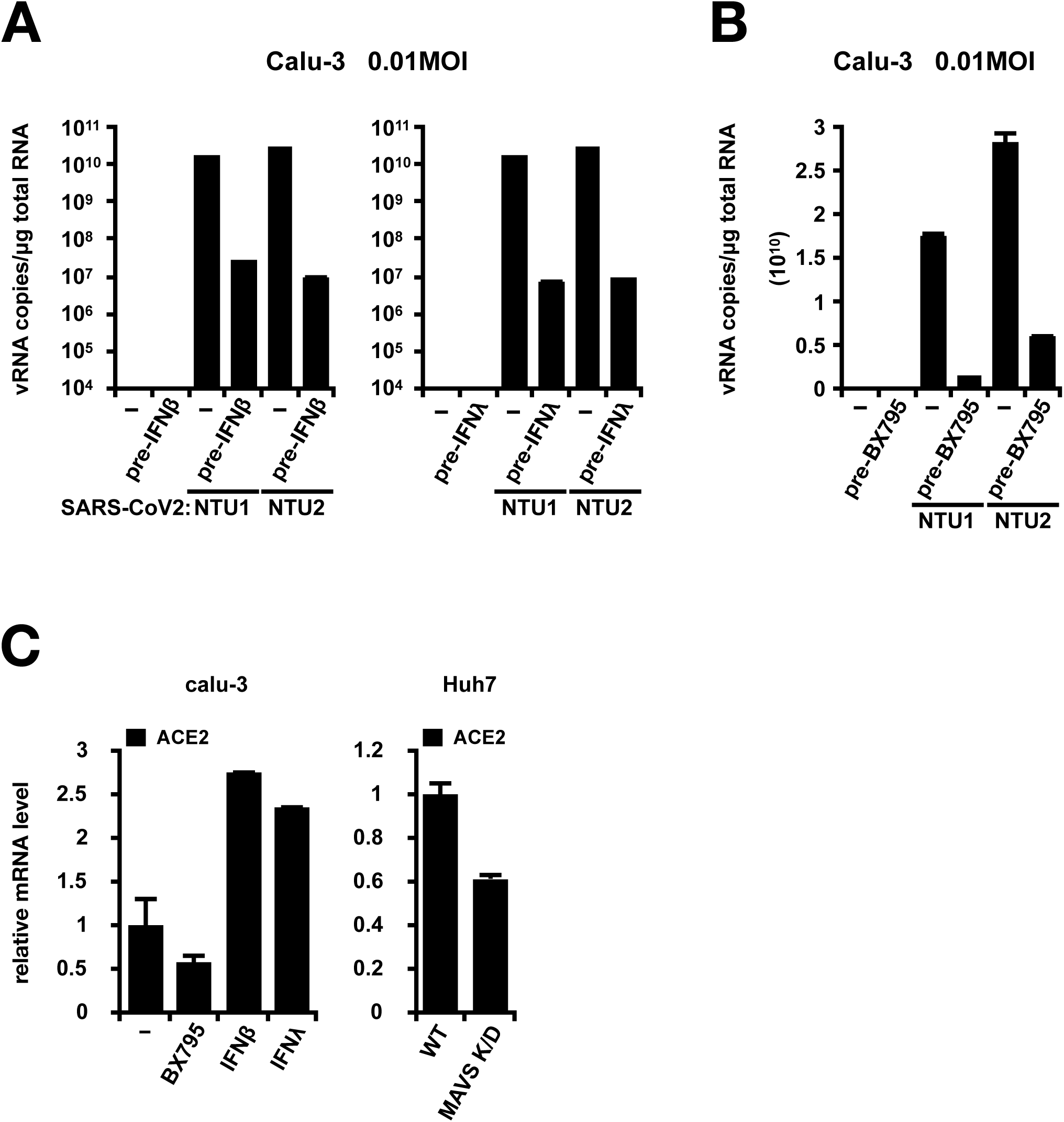
The intracellular vRNA levels of SARS-CoV2 in Calu-3 cells pretreated with IFNβ, IFNλ1 (A), or BX795 (B). Treatment of IFNβ and IFNλ1prior to SARS-CoV2 infection strongly reduced the intracellular vRNA levels detected at 24 h.p.i..(C) The mRNA expression levels of ACE2 in Calu-3 cells pretreated with IFNβ, IFNλ1and in wildtype Huh7 or MAVS K/D Huh7 cells.

**Supplementary Fig. 4.**
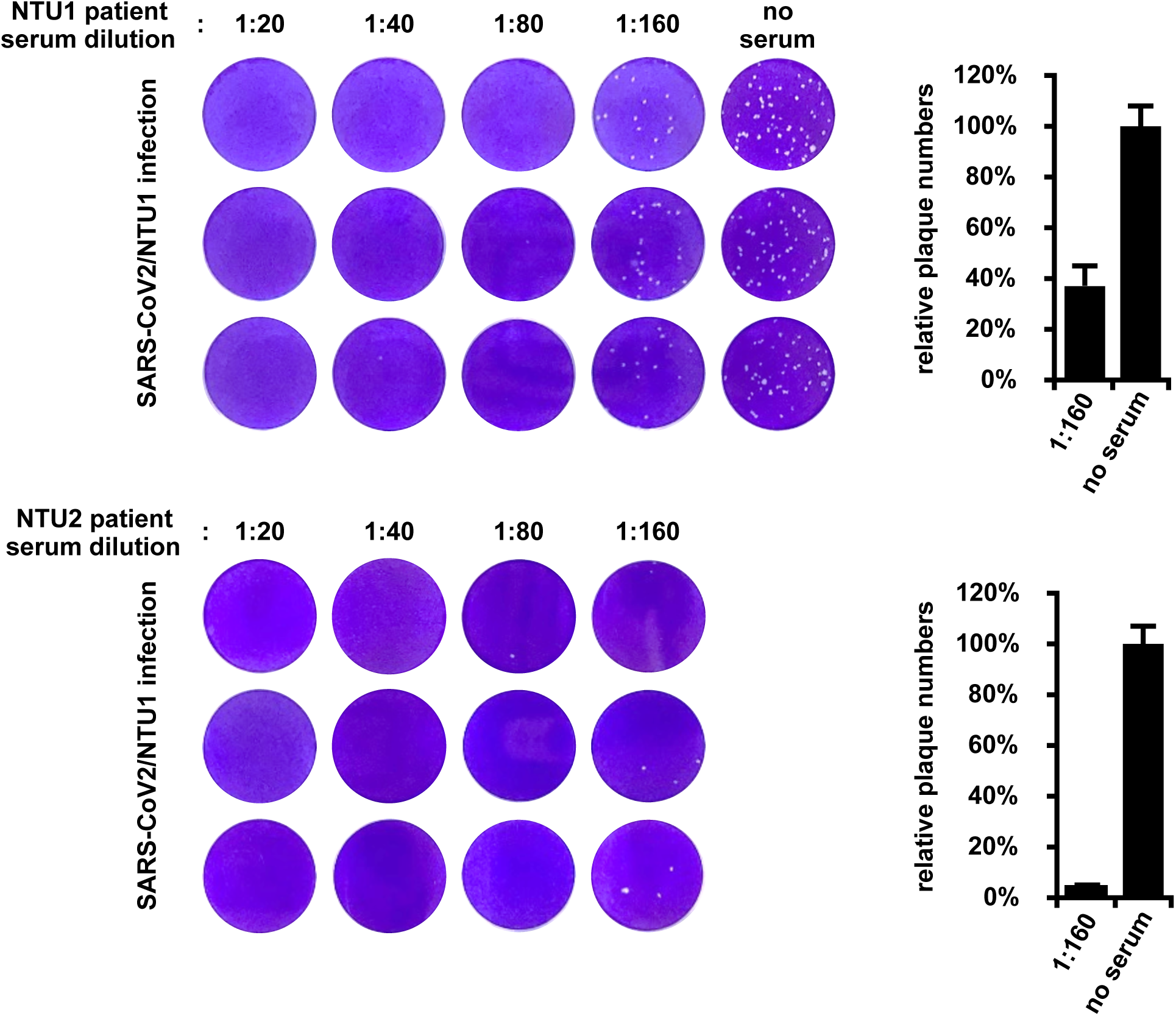
Plaque reduction neutralization of NTU1 isolate by either NTU1 or NTU2 patient serum collected post seroconversion.

**Supplementary Table 1.**
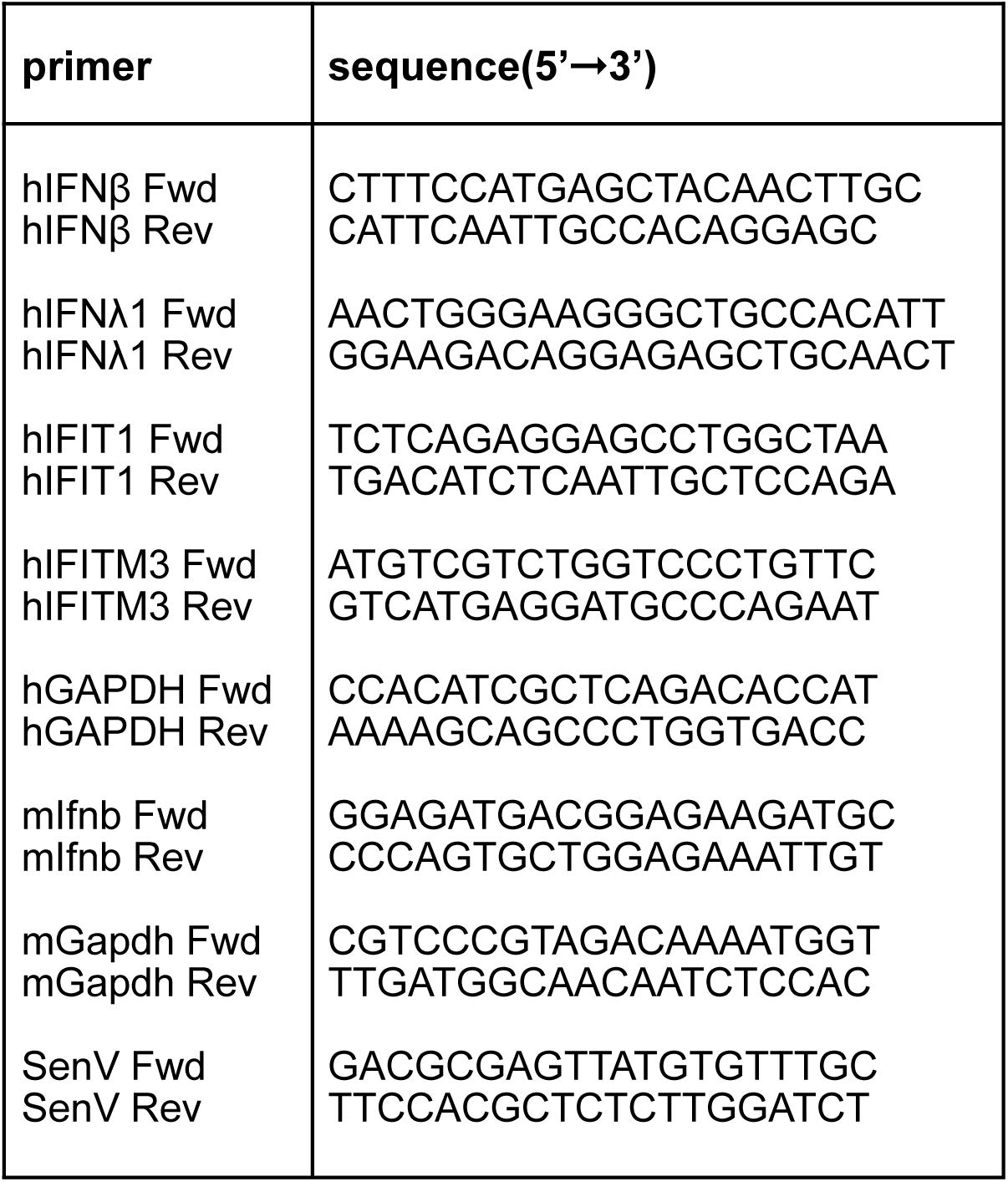

